# COSMIC-Linked Ras Mutations at the Interface Between H-Ras and PI3Kγ_RBD_ Frequently Generate Affinity Increases

**DOI:** 10.64898/2026.05.01.722339

**Authors:** Elizabeth H. Mead, Kaeden C. Batz, Kuo-Hsien Shih, Ian R. Fleming, Corey D. Tesdahl, Lucas Lizardos, Joy R. Armendariz, Jonathan P. Hannan, Ava M. Hickey, Alan Leyk, Annette H. Erbse, Joseph J. Falke

**Affiliations:** Department of Biochemistry and Molecular Biophysics Program, University of Colorado at Boulder, Boulder, CO 80309-0596, USA

## Abstract

The three conventional isoforms of the Ras G-protein (H-, K-, N-Ras) function as molecular on-off switches that regulate a wide array of signaling pathways, including the Ras-PI3K-PIP_3_-PDK1-AKT pathway that is central to innate immunity and normal cell growth, and is dysregulated in many disease states. Activation of the pathway by Ras requires adequate Ras-PI3K binding affinity. Here we focus on the interface of known structure in the H-Ras:PI3Kγ co-complex essential to multiple pathways including directed pseudopod growth in leukocyte chemotaxis. At this interface 10 H-Ras residues, all 100% conserved between the H-, K- and N-Ras isomers, contact the Ras binding domain of PI3Kγ (PI3Kγ_RBD_). To investigate the degree to which the native H-Ras:PI3Kγ_RBD_ interface is optimized by evolution for maximal binding affinity, 8 interfacial Ras mutations selected from the COSMIC database and the literature were introduced at the contact positions. All 8 Ras mutations were observed to alter the H-Ras:PI3Kγ_RBD_ binding affinity, with 4 mutations yielding significant affinity increases and 4 yielding significant affinity decreases. These findings indicate that the native H-Ras:PI3Kγ_RBD_ interface provides intermediate, rather than maximal, binding affinity. Such intermediate affinity is consistent with the substantial binding plasticity of the conserved H-, N-, K-Ras effector docking surface, which has evolved to bind a diverse array of effectors. Furthermore, the findings provide evidence that COSMIC-linked mutations at the H-Ras:PI3Kγ_RBD_ interface frequently generate affinity increases as well as decreases, with potential implications for molecular mechanisms of disease and for tool development in cell biology.

---

Activation of the PI3K-PIP_3_-PDK-AKT pathway begins with G protein and receptor signals that activate Class 1 PI3K lipid kinases, yielding production of the growth signal lipid PIP_3_^1–12^. The present study focuses on the semi-conserved, regulatory protein-protein interface formed in many pathways between an isoform of the G protein Ras (H,K,N) and a PI3K isoform (α,γ,δ), yielding a heterodimeric Ras:PI3K complex. Each Ras:PI3K interface is dominated by contacts between Ras and the structurally independent Ras binding domain (RBD) of the PI3K isoform. Ras binding to PI3K synergizes with tyrosine kinase signals (for PI3Kα and PI3Kδ), or with G_βγ_ signals (for PI3Kγ) to generate maximal lipid kinase activation and PIP_3_ production. The ensuing PIP_3_ growth signal upregulates one or more essential cell pathways, while excessive PIP_3_ production yields many pathologies.

Most studies of disease-linked Ras mutations have focused on the hotspot positions G12, G13, or Q61 within the GTPase active site that disrupt GTP hydrolysis, yielding global activation of Ras signaling pathways^1–4^. Here we investigate Ras mutations distal to the active site, focusing primarily on the 10 H-Ras positions that contact the Ras binding domain of PI3Kγ in the known structure (PDB 1HE8) of the H-Ras:PI3Kγ co-complex^12^ **(Figure 1)**. These 10 H-Ras positions lie within the Ras effector lobe (residues 1-86) where conventional Ras isoforms (H,K,N) share 100% sequence identity^1–4^. As a result, a given Ras effector protein can often bind all three Ras isoforms. Moreover, each Ras isoform binds multiple effectors with overlapping docking footprints on Ras, which presents a recognition challenge since effectors exhibit only moderate conservation of their Ras binding surfaces. Thus, the diversity of Ras effectors requires the perfectly conserved Ras effector docking surface to possess considerable binding plasticity. We hypothesize that the Ras effector docking surface is unlikely to be optimized for maximal affinity towards one effector, predicting that Ras mutations at effector contact positions could either increase or decrease effector affinity.

**Figure 1.**
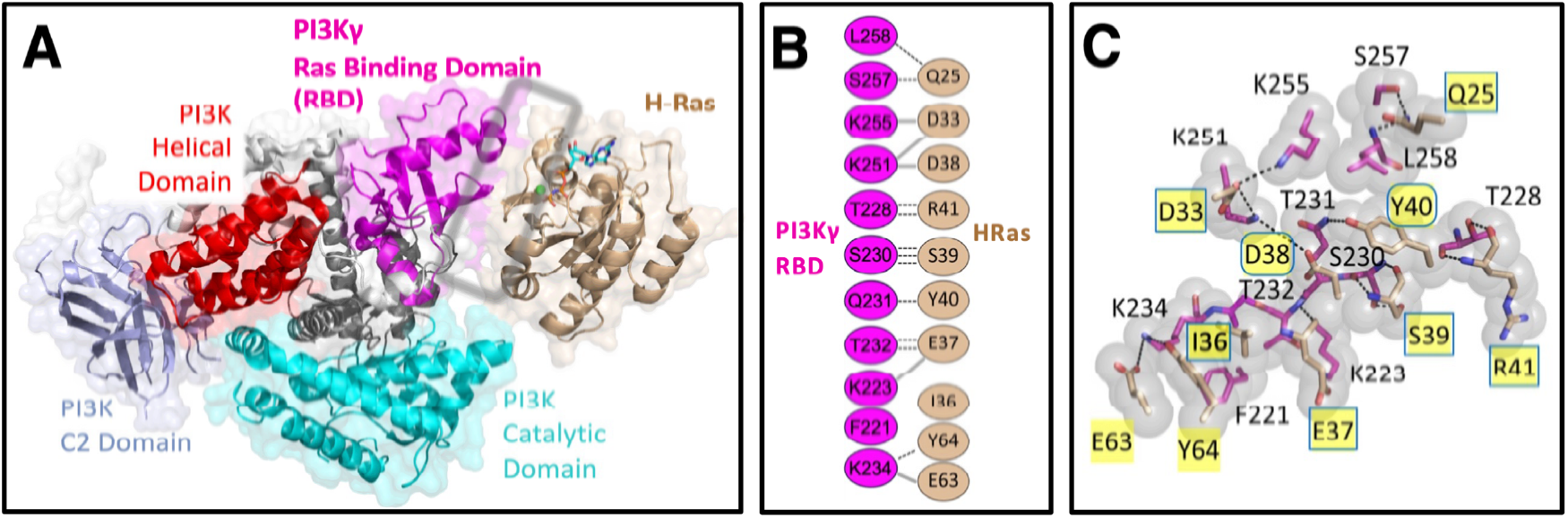
The H-Ras:PI3Kγ interface. **(A)** Crystal structure of the H-Ras:PI3Kγ co-complex (PDB 1HE8) highlighting the primary interface between H-Ras and the PI3Kγ Ras-binding-domain (PI3Kγ_RBD_)^12^. **(B)** The 10 H-Ras positions that contact PI3Kγ_RBD_ in the crystal structure exhibit 100% identity between conventional H-, N-, and K-Ras isoforms, with representative salt bridges (solid) and H-bonds (dashed)^12^. **(C)** A slice through the co-complex interface highlighting in yellow the 10 interfacial H-Ras positions that contact PI3Kγ_RBD_. Boxes and ovals specify the 8 H-Ras positions mutated in our studies, sampling most regions of the PI3Kγ_RBD_ contact surface^13, 14^.

The present study tests this prediction by introducing Ras mutations at the H-Ras:PI3Kγ_RBD_ interface and measuring their effects on the affinity of H-Ras binding to PI3Kγ_RBD_. Random, interfacial mutations at protein-protein interfaces usually weaken binding affinity^15–17^. To increase the likelihood of finding affinity increasing mutations at the H-Ras:PI3Kγ_RBD_ interface we searched the COSMIC cancer somatic mutation database^18^ for mutations in H-Ras, K-Ras and N-Ras at the 10 Ras positions that contact RBD in the H-Ras:PI3Kγ structure. We reasoned that (i) some or all of these COSMIC mutations may increase the binding affinity of one or more Ras:PI3K isoform pairs, thereby potentially supporting cancer by increasing increasing PIP_3_ growth signal production, and (ii) interfacial COSMIC mutations in all three Ras isoforms are valid targets since a mutation in any one isoform would likely perturb the identical effector docking subdomains of all three isoforms in a similar fashion. The resulting COSMIC database search yielded altogether 30+ Ras mutations at the 10 target positions of H-, K- and N-Ras. For the present study we chose 8 mutations (Q25L, D33E, I36V, E37K, S39Y, R41K, R41L, R41Q) that sample most of the H-Ras:PI3Kγ_RBD_ interface (Figure 1C). (Positions E63 and Y64 were exempted since mutations at these positions have been found to inhibit intrinsic GTPase activity, presumably due to their proximity to G12 and the GTPase site^19^).

The 8 selected mutations were introduced into a well-characterized H-Ras expression construct by site-specific mutagenesis and confirmed by plasmid sequencing. Subsequently, each mutant H-Ras protein was expressed in *E. coli*, purified, loaded with the desired nucleotide (the non-hydrolyzable GTP analogue GMPPNP, GTP, or GDP) and analyzed utilizing our published methods^13, 14, 19^ (Supporting Information) to generate an H-Ras stock of known protein concentration and fractional activation. Similarly, we used our published methods to express and characterize a construct containing the functional PI3Kγ Ras binding domain^13, 14^ (Fig. 2 and Supporting Information), yielding an RBD stock of known concentration. The resulting H-Ras and RBD stocks were subsequently employed in H-Ras:PI3Kγ_RBD_ binding titrations.

**Figure 2.**
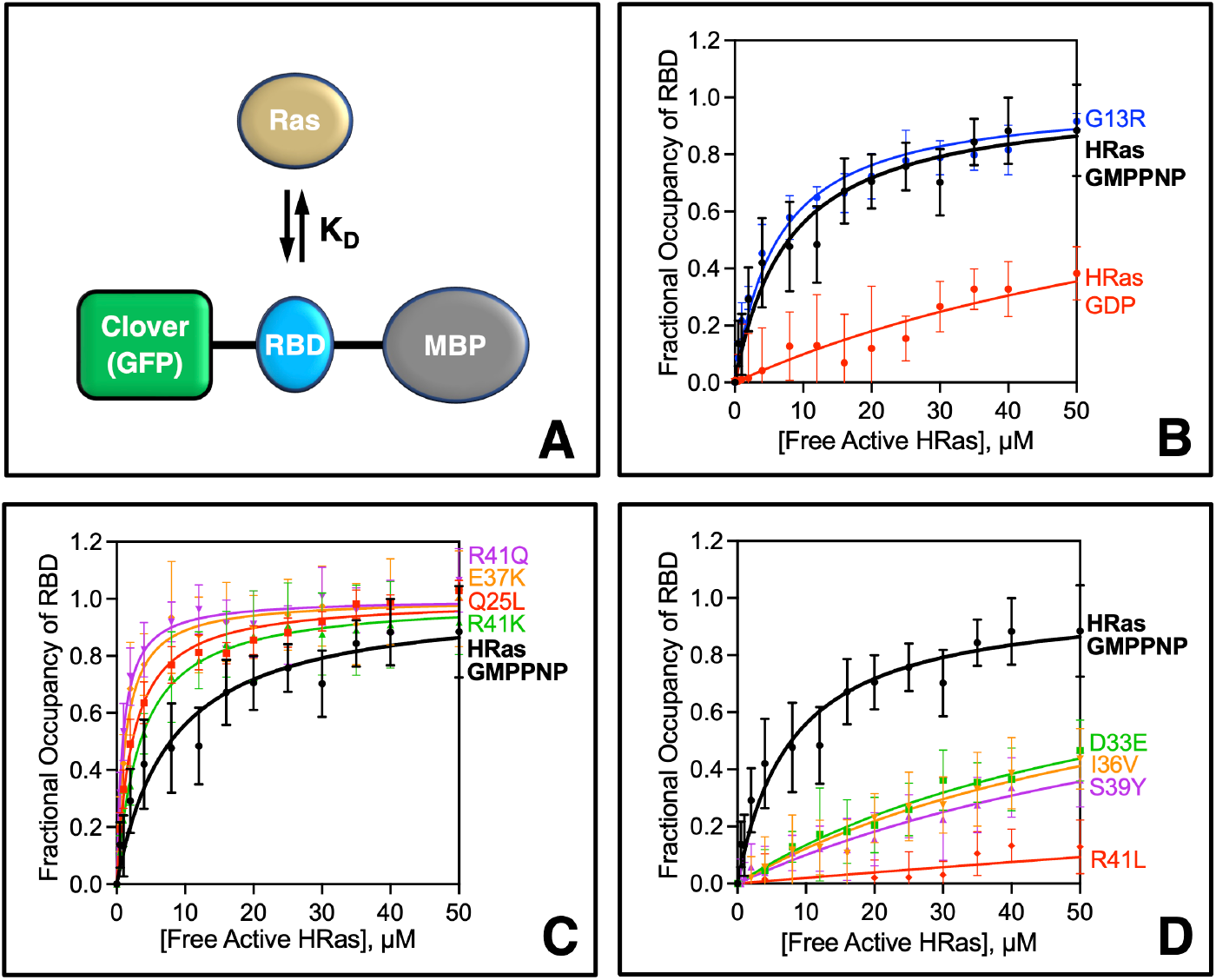
Effects of interfacial Ras mutations on H-Ras binding to PI3Kγ Ras binding domain. **(A)** Schematic binding reaction measures the equilibrium dissociation constant (K_D_) for H-Ras binding to the PI3Kγ Ras binding domain (PI3Kγ_RBD_) in a fusion construct with N-terminal maltose binding protein for stability and C-terminal Clover GFP for fluorescence^13, 14^. **(B-D)** Mean equilibrium binding curves, measured by microscale thermophoresis, quantify the affinity of the indicated H-Ras proteins for PI3Kγ_RBD_. The WT H-Ras construct possesses native residues at all 10 positions that contact PI3Kγ_RBD_, and its binding curve is displayed in each plot for comparison. Variants were created by site directed mutagenesis in the same background. All proteins loaded with activating GMPPNP unless noted otherwise. **(B)** A comparison of WT H-Ras loaded with GMPPNP and GDP confirms these two nucleotides activate and inactivate Ras binding to PI3Kγ_RBD_, respectively. Control mutant H-Ras G13R loaded with GMPPNP yielded WT binding within error, consistent with its distal location relative to PI3Kγ _RBD_ ^12^. **(C**,**D)** Interfacial mutations that significantly increase or decrease, respectively, H-Ras binding affinity for PI3Kγ_RBD_. (B-D) Error bars are ± SD; see Table 1 for N and best fit K_D_ values. Each binding curve averaged at least 8 triplicate titrations carried out on different days by at least 2 independent researchers using at least 2 independent protein preparations. Detailed methods in Supporting Information.

Microscale thermophoresis (MST) binding titrations were carried out to quantify the affinity of each H-Ras mutant for the isolated Ras-binding-domain of PI3Kγ utilizing our validated and published protocol^14^ (**Figure 2** and Methods, Supporting Information). Each titration employed 16 samples containing a fixed concentration of the PI3Kγ_RBD_ construct (100 nM) and increasing concentrations of active H-Ras (0 to 50 µM), where active H-Ras is operationally defined as H-Ras loaded with activating nucleotide (GMPPNP or GTP). Each titration was carried out in triplicate, along with a side-by-side triplicate titration of the H-Ras Q25L mutant that we have previously shown effectively saturates the PI3Kγ_RBD_ construct at the highest H-Ras concentrations employed. The resulting average H-Ras Q25L binding curve served as an internal reproducibility standard, and its best-fit asymptote yielded the maximum MST signal at saturation used for binding curve normalization. Conditions ensured that the concentration of bound active H-Ras was always less than 5% that of total active H-Ras, such that [Total Active Ras] ≈ [Free Active Ras] within error. Figures 2C-D plot the mean binding curve for each mutant, obtained by averaging at least 8 triplicate titrations.

**Table 1.** Effects of 8 Interfacial Ras Mutations on H-Ras Binding to PI3Kγ _RBD_.

| H-Ras Protein | $K_D \pm \text{SEM}$ ($\mu\text{M}$)<sup>a</sup> | Significance (p-Value)<sup>b</sup> | Number of Triplicates | Number of Protein Preps | Normalization Factor<sup>c</sup> |
| --- | --- | --- | --- | --- | --- |
| WT H-Ras GMPPNP | $6.8 \pm 1$ | 1 | 8 | 3 | 1.0 |
| G13R (Distal) | $6.3 \pm 1$ | 0.398 | 11 | 3 | 1.0 |
| Q25L | $2.8 \pm 0.1$ | <0.001 | 90 | 17 | 1 |
| D33E | >50 $\mu\text{M}$ | <0.001 | 8 | 3 | 1.0 |
| I36V | >50 $\mu\text{M}$ | <0.022 | 9 | 3 | 1.0 |
| E37K | $1.3 \pm 0.1$ | <0.001 | 10 | 3 | 0.8 |
| S39Y | >50 $\mu\text{M}$ | <0.001 | 11 | 2 | 1.0 |
| R41K | $3.5 \pm 0.2$ | <0.001 | 9 | 3 | 1.5 |
| R41L | >50 $\mu\text{M}$ | <0.014 | 8 | 2 | 1.0 |
| R41Q | $1.0 \pm 0.2$ | <0.001 | 8 | 3 | 1.1 |
| WT H-Ras GDP | >50 $\mu\text{M}$ | <0.001 | 8 | 2 | 1.0 |

<sup>a</sup> $K_D$ values were measured for the indicated H-Ras mutants binding to PI3Kγ <sub>RBD</sub>.
<sup>b</sup> Two-tailed t-tests yielded the indicated p values comparing the $K_D$ of each H-Ras mutant to that of WT H-Ras GMPPNP possessing native interfacial residues.
<sup>c</sup> Normalized relative to the saturating MST signal of H-Ras Q25L, defined by the best fit asymptote of the H-Ras Q25L binding curve. A value of 1.0 indicates that the maximum MST signal of the mutant was indistinguishable from that of the internal standard H-Ras Q25L. A value differing from 1.0 indicates the maximum MST signal of the mutant differed from that of H-Ras Q25L (Methods in Supporting Information).

**Table 1** summarizes the resulting best-fit equilibrium dissociation constants (K_D_) measured for the binding of each H-Ras variant to PI3Kγ_RBD_. Notably, the WT H-Ras construct that possesses all 10 native contact residues yielded K_D_ = 6.8 ± 1 µM for binding to PI3Kγ_RBD_, a value within 2-fold of that observed for H-Ras binding to full length PI3Kγ^12, 14^. This suggests that H-Ras binding to RBD dominates the native binding interaction, and that the RBD model system is useful for analyzing the effects of mutations on binding affinity. Four interfacial Ras mutations (Q25L, E37K, R41K, R41Q) significantly increased the affinity of Ras binding to PI3Kγ_RBD_ between 2.0- and 6.8-fold relative to the WT Ras construct (Figure 2C, each p < 0.001). In contrast, 4 other interfacial Ras mutations (D33E, I36V, S39Y, R41L) significantly decreased the Ras-PI3K_RBD_ binding affinity at least 5-fold (Figure 2D, p < 0.001 to p < 0.022). The latter four low-affinity binding curves did not approach saturation; as a result, their K_D_ values were not well defined but the maximum free active Ras concentration employed in the titration provides a lower limit K_D_ > 50 µM in each case.

The results indicate that Ras mutations at the H-Ras:PI3Kγ_RBD_ interface can yield both affinity increases and decreases. It follows that the interface is sensitive to mutational perturbation at multiple contact positions and has not been optimized by evolution to provide maximal binding affinity for a representative effector. Instead, the H-Ras:PI3Kγ_RBD_ interface has evolved an intermediate affinity that is adequate for the biological function of H-Ras:PI3Kγ regulation in cellular pathways. Such intermediate affinity is consistent with the plasticity needed for the conserved Ras effector docking surface to bind with adequate affinity to diverse effectors. At the same time, intermediate affinity makes this surface susceptible to mutations that increase affinity in disease or in tool development.

The observation that 8 COSMIC-linked Ras mutations at the H-Ras:PI3KγRBD interface yield 4 affinity increases and 4 affinity decreases supports the hypothesis that COSMIC mutations at this interface exhibit similar propensities for affinity increases and decreases. This observation differs from the strong preponderance of affinity decreases typically observed for random mutations at protein-protein interfaces. Current large databases of affinity changes generated by random interfacial mutations in a diverse set of binding partners display an approximately 1:4 ratio of affinity increases to decreases^17^. Simple coin flip statistics^20, 21^ for a 1:4 ratio of likelihoods yields only a 6% probability that 8 mutations will yield 4 or more affinity increases. Thus the tested COSMIC mutations exhibit a considerably greater frequency of affinity increases than expected for typical random interfacial mutations. When the ratio of likelihoods is 1:1, the coin flip probability of 8 mutations yielding 4 or more affinity increases rises to 63%. Although oversimplified, this analysis is consistent with the hypothesis that COSMIC-linked Ras mutations at the H-Ras:PI3KγRBD interface (a) generate affinity increases and decreases with similar frequencies, and (b) generate a higher ratio of affinity increases to affinity decreases than random point mutations at typical interfaces.

The affinity effects of specific Ras interfacial mutations may well differ among Ras:PI3K isoform pairs. Unlike the identical effector binding subdomains of H-, K- and N-Ras, the Ras binding domains of PI3Kα, γ and δ are moderately homologous rather than identical. For example, a comparison of the recently determined structure of the K-Ras:PI3Kα:GlueD927 co-complex (PDB 9C15)^22^ with that of the H-Ras:PI3Kγ co-complex (PDB 1HE8)^12^ reveals their Ras:PI3K_RBD_ interfaces possess both similarities and differences. The latter study investigated the effects of interfacial Ras mutations on K-Ras:PI3Kα_RBD_ binding affinity, providing information complementary to the present findings for the H-Ras:PI3Kγ_RBD_ interface. Table S1 (Supplementary Information) summarizes (i) the effect on Ras:PI3K_RBD_ binding affinity, (ii) the COSMIC status, and (iii) the number of COSMIC hits in H-, K-, and N-Ras for each interfacial Ras mutation in both studies. Altogether, the table includes the 8 COSMIC-linked interfacial Ras mutations described herein (Table 1) and 2 additional COSMIC-linked Ras mutations at the K-Ras:PI3Kα_RBD_ interface (the latter study focused primarily on alanine scanning rather than on COSMIC mutations). The 2 COSMIC-linked Ras mutations at the K-Ras:PI3Kα_RBD_ interface yield 1 significant, 3.3-fold affinity increase and 1 no change, within error. These additional independent findings, though small in number, remain consistent with the hypothesis that a substantial fraction of COSMIC-linked Ras mutations trigger affinity increases at Ras:PI3K_RBD_ interfaces.

Tumor-linked mutations in cancer databases can be classified as high impact driver mutations that play central roles in tumorigenesis, medium impact mutations that support drivers, and low-impact passenger mutations^23^. Ras driver mutations are found at Ras positions 12, 13 and 61^1–4^. It is plausible to hypothesize that interfacial COSMIC-linked mutations found to significantly increase Ras:PI3K binding affinity may represent medium impact mutations capable of supporting drivers by stimulating Ras-PI3K-PIP_3_ signaling. Multiple lines of evidence indicate that membrane-anchored Ras activates PI3Ks primarily by direct binding and membrane recruitment^11, 24–28^. Thus, in the simplest scenario, the 2.0-to 6.8-fold affinity increases observed for the affinity-enhancing mutations would yield a proportional increase in PI3K binding, membrane density and net PIP_3_ production. Activation of PI3K at these levels has been shown to be oncogenic, for example the most oncogenic PI3K mutation, PI3Kα H1047R, drives tumorigenesis by generating 3-to 7-fold increases in PI3Kα membrane affinity, density, and net production of PIP_3_ signaling lipid^24, 28^. At least two features of the Ras-PI3K-PIP_3_-PDK-AKT pathway explain how such modest changes in PIP_3_ levels can generate large changes in downstream growth signals^1–4, 29^: (i) PDK1 phospho-activation of AKT1 is expected to exhibit a quadratic dependence on PIP_3_ density since the reaction requires simultaneous recruitment of PDK1 and AKT1 to the target membrane via binding of their PH domains to independent PIP_3_ molecules; and (ii) the pathway is a multi-step, self-amplifying, exponential cascade in which the product molecules of a given step each generate multiple product molecules in the next step. Interestingly, COSMIC links the affinity-enhancing mutations in Table S1 to 6 H-Ras associated cancers, 4 K-Ras cancers and 1 N-Ras cancer. While small numbers, the apparent dominance of COSMIC-linked H-Ras and K-Ras affinity-enhancing mutations is consistent with the established roles of H-Ras:PI3Kγ signals in innate immunity and cancer, and of K-Ras:PI3Kα signals in cell growth and cancer^1–9^.

The 8 COSMIC-linked, interfacial H-Ras mutations found herein to perturb H-Ras:PI3Kγ_RBD_ binding affinity may also alter Ras binding to other effectors. The docking footprints of multiple Ras effectors, including B-Raf as well as PI3K isoforms α,γ,δ ^22, 25, 30, 31^, exhibit partial overlap on the conserved effector docking face of Ras. Thus, in principle, a given COSMIC-linked interfacial Ras mutation could impact a single pathway, or multiple pathways. The present results for H-Ras R41Q show this mutation confers increased H-Ras:PI3Kγ_RBD_ binding affinity, which is predicted to yield increased activation of Ras-PI3K-PIP_3_-PDK-AKT signaling in cells. Previous studies found this mutation has an inhibitory effect on the Ras-Raf-MEK-ERK growth signal pathway in cells^32^. It follows that R41Q may be an example of a medium impact mutation that appears in COSMIC because it selectively activates the Ras-PI3K-PIP_3_-PDK-AKT pathway. In contrast, E37K may upregulate both major growth pathways since it increases affinity for PI3K_RBD_ (Table 1) and increases Ras-Raf-MEK-ERK signaling in cells^33–35^. In pathways controlled by different Ras:PI3K isoform pairs, interfacial Ras mutations may have contrasting effects on different PI3K isoforms due to their non-identical Ras binding domains. For example, the co-complex interfaces of H-Ras:PI3Kγ^12^ and K-Ras:PI3Kα:GlueD927^22^ share 9 identical Ras residues that contact PI3K_RBD_ in both structures, but also exhibit 1 unique Ras contact residue and 4 unique Ras contact residues, respectively. Moreover, Ras and PI3K isoforms exhibit differential tissue expression, and even in the same cell can target to different regions of the plasma membrane. In short, the biological impacts of a given Ras mutation may vary between tissues, membrane locations, pathways, and specific effectors^36, 37^.

Additional studies are needed to test key hypotheses arising from the present findings. The prediction that COSMIC-linked interfacial Ras mutations generate Ras-PI3K_RBD_ affinity increases more often than random mutations can be tested by comparing the affinity impacts of interfacial Ras mutations listed in COSMIC to those of random, non-COSMIC linked mutations. The prediction that the Ras-PI3K_RBD_ affinity increases observed for disease-linked interfacial Ras mutations will yield proportional increases in Ras-PI3K affinity, PI3K membrane recruitment and net PIP_3_ growth signal production can be tested by biophysical studies of Ras-PI3K-PIP_3_ signaling reactions employing full length proteins reconstituted on a target membrane surface^11^. Such studies are also well suited for comparing the effects of interfacial Ras mutations in different Ras:PI3K isoform pairs. More broadly, it is important to identify the biological impacts of interfacial, affinity-changing Ras mutations via cell and animal studies of Ras-activated pathways including Ras-PI3K-PIP_3_-PDK-AKT and Ras-Raf-MEK-ERK. Of special interest would be the discovery of interfacial mutations that specifically activate or inhibit only a single pathway, which could serve as research tools for up- or down-regulation of specific Ras effector pathways in live cells. A testable prediction is that a Ras double mutant possessing both a hotspot driver mutation to stabilize the on-state and a pathway-specific, affinity-altering mutation could be a pathway super-driver or blocker. Finally, molecular dynamics simulations of the effects of disease-linked Ras mutations on Ras:effector interfaces will provide a deeper understanding of these interfaces and the forces that stabilize them.

## Supporting information

Supporting Information - Methods, Fig S1, Table S1, Contributions

## Acknowledgements

The authors are grateful for funding by NIH/NIGMS Grant R35GM144346 to JJF and Traineeship T32GM142607 to JA; Beckman Scholar Award to IF; Wuttke/Beckman Biomedical Sciences Research Award to AH; CU Boulder UROP Awards to KB and CT. We thank the Biochemistry Shared Instruments Pool (RRID SCR_018986) for the use of centrifuges (NIH Grant R24OD033699) and MST (NIH Grant S10D21603).

## Supporting Information

Contents include 1) Methods, 2) Figure S1 illustrating UV deconvolution and HPLC traces methods quantifying H-Ras concentration and nucleotide loading, 3) Table S1 providing data for Ras mutations at the H-Ras:PI3Kγ_RBD_ and K-Ras:PI3Kα_RBD_ interfaces including affinity changes, COSMIC hits in specific Ras isoforms, nucleotide substitutions, and 4) Author Contributions.

## Accession Identifiers for Proteins

### WT H-Ras

The WT H-Ras construct employed herein is a derivative of human H-Ras (UniProt Identifier P01112) with the modifications specified in Methods (Supporting Information).

### PI3Kγ Ras Binding Domain (RBD)

The RBD construct employed herein is a fragment of human PI3Kγ (Uniprot Identifier P48736) as specified in Methods (Supporting Information).

