## Supporting Information - Methods, Fig S1, Table S1, Contributions for "COSMIC-Linked Ras Mutations at the Interface Between H-Ras and PI3Kγ_RBD_ Frequently Generate Affinity Increases"

University of Colorado at Boulder

Boulder, CO 80309-0596

USA

* To whom correspondence should be addressed:

### CONTENTS:

**1) Detailed Methods, pages S.2-9**

**2) Figure S1, page S.9,** "Representative UV Deconvolution and HPLC Analyses Used to Quantify the Concentration and Nucleotide Loading of H-Ras Stocks for Binding Measurements"

**3) Table S1, page S.10,** "Effects of 20 Interfacial Ras Mutations on Ras Binding to PI3K$_{RBD}$ as Measured for the H-Ras:PI3K$\gamma_{RBD}$ and K-Ras:PI3K$\alpha_{RBD}$ Isoform Pairs"

**4) Contributions, page S.11**

**5) References for Methods, page S.11-12**

### METHODS

**Reagents**. *E. coli* BL21 cells were obtained from Thermo Fisher Scientific (Waltham, MA), and Bacto yeast extract and tryptone for bacterial growth were from Gibco (Waltham, MA). Granulated agar for bacterial plates was from Becton Dickinson (Sparks, MD). Ampicillin, HEPES free acid, and phenylmethyl sulfonyl fluoride (PMSF) were from Research Products International (Mount Prospect, IL). Reduced L-glutathione and ethylenediamine tetracetic acid disodium salt dihydrate (EDTA) were from ThermoFischer Scientific (Waltham, MA). Kanamycin, bovine serum albumin (BSA), and Tween 20 were obtained from Sigma-Aldrich (St. Louis, MO). Talon cobalt(II) metal affinity resin for polyhistidine tag purification was from Takara (Kusatsu, Japan), and imidazole was from Electron Microscopy Sciences (Hatfield, PA). Non-hydrolysable GTP analog GMPPNP, conjugated with four lithium counter ions, at >95% purity, was from Abcam (Cambridge, UK). Guanosine 5'-diphosphate [GDP] disodium salt was from Sigma-Aldrich (St. Louis, MO). Desalting columns were Econo-Pac 10DG desalting prepacked gravity flow columns from Bio-Rad (Hercules, CA). Vivaspin 500 spin concentrators, 10,000 MWCO from Sartorius were used to concentrate protein samples (Göttingen, Germany). Microscale thermophoresis measurements were performed on a Nanotemper Monolith NT.115 instrument utilizing Monolith NT.115 capillaries (Munich, Germany).

**H-Ras Plasmid Construct and Mutagenesis.** Our previously published H-Ras expression construct, a modified version of an H-Ras construct kindly gifted by the Groves laboratory, was employed as the background for the new mutants described herein[1-4]. This H-Ras C118S/C181S Δ185-189 construct is otherwise identical the wild-type H-Ras sequence and retains all native contact residues for PI3Kγ Ras binding domain, and is referred to as "WT" H-Ras. The other H-Ras variants employed were created in this H-Ras background by introducing a single point mutation (G13R, Q25L, D33E, I36V, E37K, S39Y, R41K, R41L and R41Q) using the QuikChange mutagenesis kit from Agilent Technologies (Santa Clara, CA) and mutagenesis DNA primers from Integrated DNA Technologies (Coralville, IA), followed by sequencing of the full H-Ras gene by GENEWIZ (South Plainfield, NJ) to confirm the desired mutation and that no unintended mutations were present. To isolate plasmid for sequencing and future protein expression, the mutagenesis reaction was transformed into DH5α *Escherichia coli* cells and grown on 2xYT agar plates containing 50 μg/mL ampicillin for 18 hours at 37°C. Single colonies were inoculated into culture flasks with 5 mL of 2xYT media and the same ampicillin concentration. Following 15 hours of shaking incubation at 37°C, the OMEGA Bio-Tek E.Z.N.A. Plasmid DNA Mini Kit (Kansas City, MO) was used to purify the H-Ras plasmid.

**Ras Protein Expression and Purification.** Recombinant H-Ras expression and purification was carried out as we previously described with the following minor modifications[2-4]. Briefly, H-Ras plasmids were transformed in BL21(DE3) chemically competent *E. coli* cells and grown on 2xYT + agar plates with 100 μg/mL ampicillin for 16 hours at 37°C. Single colonies were inoculated into 12.5 mL of 2xYT media supplemented with 50 μg/mL ampicillin and grown for 15 hours at 37°C, shaking at 250 rpm. This culture (12.5 ml) was diluted into 750 mL of media of the same composition and grown as above for approximately 2.5-3 hours until an OD600 reading of 0.4 was reached. Protein expression was induced by addition of isopropyl β-D-1-thiogalactopyranoside (IPTG) (Invitrogen, Waltham MA) to 500 μM then the cultures were incubated with shaking at 18°C for 18 hours. Cells were pelleted by centrifugation at 4,000 x g for 15 minutes at 4°C and then resuspended to 35 mL total volume in ice cold Lysis Buffer (50 mM $Na_2HPO_4$, 150 mM NaCl, 1 mM $MgCl_2$, 2.5mM glutathione, 1 mM PMSF, pH 8.0). Cells were lysed by sonication and cellular debris were removed by centrifugation at 30,000 x g for 25 minutes at 4°C. The clarified supernatant was passed through a 20 mL Bio-Rad Econo-Pac® chromatography column (Hercules, CA) containing 1.5 mL Talon cobalt(II) resin and the 6X N-terminal His-tagged Ras protein bound to the resin was successively washed with 9 mL of ice cold Wash 1 buffer (50 mM HEPES, 150 mM NaCl, 1 mM $MgCl_2$, 2.5mM glutathione (fresh), 1 mM PMSF, pH 8.0) and 6 mL of ice cold Wash 2 Buffer (Wash 1 Buffer + 10mM imidazole), then eluted with 4 mL of Elution Buffer (50 mM HEPES, 300 mM NaCl, 1 mM MgCl2, 250 mM imidazole, 2.5mM glutathione, pH 8.0) all in a cold room at 4°C.

**Ras Nucleotide Loading, Protein Concentration, and Storage.** To load the purified H-Ras with GTP, non-hydrolysable analogue GMPPNP, or GDP our previously described procedure was used, with the following minor modifications[2-4]. Briefly, eluted protein samples were transferred into Exchange Buffer (25mM HEPES, 140mM KCl, 15mM NaCl, 2.5mM glutathione, pH 7.4) using Bio-Rad desalting columns at 4°C. Subsequently, PhosSTOP (Sigma-Aldrich, Waltham, MA), a blend of phosphatase inhibitors, was added to the concentration recommended by the manufacturer and incubated at 37°C for 20 minutes to eliminate phosphatase activity. Exchange of endogenous nucleotide bound to Ras for a desired loading nucleotide was carried out at 37°C by the following modification our previous protocol. First, Ras-bound $Mg^{2+}$ was chelated via addition of 10X EDTA stock buffer (100 mM EDTA in 25 mM HEPES, 140 mM KCl, 15 mM NaCl, pH 8.0) at 1:10 dilution to yield 10 mM EDTA final. Then, nucleotide exchange was triggered by addition of either activating GMPPNP nucleotide (a non-hydrolysable analogue of GTP), activating GTP or inactivating GDP nucleotide from a 200 mM stock in exchange buffer to bring the exogenous nucleotide concentration to 30x the concentration of Ras. The ~1:10 mole ratio of $Mg^{2+}$ to EDTA employed during the nucleotide exchange reaction yielded significant $Mg^{2+}$ chelation, enabling adequate removal of bound $Mg^{2+}$ from H-Ras to speed its nucleotide exchange, while minimizing the protein destabilization that could result from more extreme levels of $Mg^{2+}$ chelation. Following the 20 min exchange reaction at 37°C, excess $Mg^{2+}$ was added via 1:10 dilution of 10X $Mg^{2+}$ buffer (110 mM $MgCl_2$, 250 mM HEPES, 140 mM KCl, 15mM NaCl, pH 8.0) to restore Ras to its native $Mg^{2+}$-bound state with very slow rates of nucleotide dissociation and exchange. The resulting sample was run through a Bio-Rad desalting column to remove unbound nucleotide while exchanging the protein-nucleotide complex into final Storage Buffer (25 mM HEPES, 140 mM KCl, 15 mM NaCl, 1 mM $MgCl_2$, 10% glycerol, 2.5 mM glutathione, pH 7.4).

After exchanging the sample of Ras-nucleotide complex into Storage Buffer its Ras protein concentration was determined by acquiring its UV spectrum then using our previously published procedure to deconvolute the protein and nucleotide components[4]. Subsequently the Ras protein concentration was calculated from the $A_{280}$ of the protein component via Beer's Law and the apo-Ras $\varepsilon_{280}$ of 19,370 $M^{-1}cm^{-1}$. Then Vivaspin 500 spin concentrators, 10,000 MWCO were used to concentrate the protein to 100-200 μM and finally a high-speed centrifugation step was performed at 372,000 x g for 20 min at 4°C to pellet aggregates before snap freezing 55 μL aliquots of supernatant in liquid nitrogen, followed by storage at -80º C.

**PI3Kγ RBD Construct: Purification, Characterization and Storage.** We previously described the expression plasmid for our PI3Kγ Ras Binding Domain construct ($PI3K\gamma_{RBD}$) employed in MST measurements of H-Ras binding to $PI3K\gamma_{RBD}$[2,5]. This construct contains (i) an N-terminal 6x His tag and maltose binding protein (MBP) region for affinity chromatography and to enhance $PI3K\gamma_{RBD}$ stability, respectively, (ii) an unstructured linker into which the functional $PI3K\gamma_{RBD}$ Ras binding domain (PI3Kγ residues 220-311) is inserted, and (iii) a C-terminal green fluorescent protein (GFP) Clover.

Our previously published protocol was employed to express and purify the $PI3K\gamma_{RBD}$ construct with the following minor modifications[2]. The expression plasmid was transformed into *E. coli*, then plated on 50 μg/mL kanamycin 2xYT agar plates. A single colony was used to inoculate 50 mL of 2xYT media containing 50 μg/mL kanamycin. Following 2 hours of incubation at 37°C with shaking at 250 rpm, 50 μM IPTG was added to induce protein expression followed by 20 hours of incubation at 30°C with shaking at 250 rpm. Cells were pelleted by centrifugation at 4,000 x g at 4°C and resuspended to 35 mL total volume in Lysis Buffer (see above). This resuspension was sonicated, and cellular debris was removed by centrifugation at 14,000 x g at 4°C. The supernatant was passed through 1.5 mL Talon cobalt(II) resin to bind to the His tag, and the bound protein was successively washed with 9 mL of ice cold Wash 1 and 6 mL of ice cold Wash 2 Buffer then eluted with 3 mL of Elution Buffer. The eluent was run through a Bio-Rad desalting column to exchange the sample into its final Storage Buffer (25 mM HEPES, 140 mM KCl, 15 mM NaCl, 1 mM $MgCl_2$, 10% glycerol, 2.5 mM glutathione, pH 7.4). The concentration of the MBP-RBD-GFP protein in each sample was determined by measuring GFP absorbance at 505 nm on a NanoDrop One spectrophotometer (ThermoFisher), followed by application of Beer's Law and the known $\varepsilon_{505}$ of Clover GFP ($\varepsilon_{505}$ = 111,000 $M^{-1}cm^{-1}$). Finally, the purified protein was adjusted with Storage Buffer to ~40 μM final concentration, then snap frozen in 30 μL aliquots in liquid $N_2$ and stored at -80°C.

**HPLC analysis of the nucleotide mixture bound to Ras, and quantification of fractional Ras activation.** Our previously published and validated HPLC procedure was used, with the following minor modifications, to quantify the mixture of guanine nucleotides (GXP) bound to H-Ras for each protein prep, yielding the fraction of the population loaded with activating nucleotide(s)[2,3]. Briefly, following nucleotide loading, WT or variant GXP:H-Ras complexes were thawed on ice then washed 5 times in Wash Buffer (25 mM HEPES, pH 7.4, 140 mM KCl, 15 mM NaCl) to remove any remaining unbound nucleotides via centrifugation in spin dialysis concentrators (Vivaspin 500 with10 kDa MW cutoff membrane (Sartorious)). Next the washed GXP:H-Ras complexes were transferred to ultracentrifuge tubes and were spun at 80,000 rpm for 10 min at 4º C in a Beckman TL-100 tabletop ultracentrifuge to eliminate nonnative aggregates. The supernatant containing washed GXP:H-Ras complexes was harvested and the bound nucleotides were isolated from protein using a heat-extraction procedure in which the washed GXP:H-Ras sample was heated at 95 °C for 6 minutes using a MiniAmp thermocycler (Applied Biosystems) to denature the protein and release its bound nucleotide. Samples were then chilled on ice for two minutes and precipitated protein was then removed by centrifugation at 11,000 rpm for 10 min at room temperature in a Beckman Microfuge 20 resulting in a clarified solution of nucleotides derived from WT or variant H-Ras. HPLC-UV was used to determine the nucleotide composition of each clarified solution using an Agilent Technology 1260/1290 Infinity HPLC system in conjunction with a Phenomenex Gemini C18 reverse-phase analytical column fitted with a Phenomenex Standard Guard Cartridge System. Mobile Phase conditions were 92.5 mM potassium phosphate ($KH_2PO_4$), 9.25 mM tetrabutylammonium bromide, pH to 6.4 with 7.5% acetonitrile. All samples were run in the mobile phase at a flow rate of 1.3 ml/min for 9 min at 22 °C. A mixture of standard nucleotides made without heating was injected to test proper HPLC system operation and confirm the characteristic retention times for each nucleotide in the UV absorbance trace (Figure S1 D), which gradually shift with normal aging of the column. In parallel the GMPPNP stock used to load the Ras prep was analyzed by HPLC without heating and always observed to be >95% pure. Next, following heat extraction from a WT or variant H-Ras protein, each clarified GXP nucleotide mixture was injected, and the identities of the individual nucleotide components were determined from the retention times of their UV peaks as illustrated for averages of triplicates in Figure S1 E-G. Note that the ΔGMPPNP component represents the GMPPNP molecules extracted from the GMPPNP-activated H-Ras subpopulation, which are intentionally and quantitatively converted by thermal hydrolysis from GMPPNP to ΔGMPPNP (or GMPPN) during the heat extraction step[3].

The individual extracted nucleotides observed for a given H-Ras sample were quantified by integration of their HPLC peaks and their relative peak areas were used to calculate the fractional activation of the starting H-Ras population as follows[3]. The H-Ras subpopulation in the active state was defined by the sum of the activating nucleotides (ΔGMPPNP (+) GTP). The H-Ras subpopulation in the inactive state was defined by the sum of the GDP-derived, inactivating nucleotides (GDP). Together these yielded the fraction of the total H-Ras population in active state, calculated as the ratio of the activating nucleotides (ΔGMPPNP (+) GTP) to the total nucleotides (ΔGMPPNP (+) GTP (+) GDP). Notably, the fractional activation of the H-Ras population was observed to vary up to 50% between different H-Ras preps, due largely to variation in the efficiency of the gentle GMPPNP loading step. This variation emphasized the importance of HPLC quantitation of each protein prep to enable accurate and reproducible titrations with active H-Ras in MST binding studies.

**Microscale Thermophoresis Measurements of Ras-RBD Binding.** The Ras-RBD MST binding assay was carried out as previously published, with the following minor modifications, to determine the equilibrium dissociation constant ($K_D$) for the binding of each H-Ras protein to the isolated RBD in solution[2]. First, Ras protein aliquots were thawed on ice and Vivaspin 500 centrifugal concentrators were rinsed with 500 µL of Ligand Buffer (25mM HEPES, 140mM KCl, 15mM NaCl, 1mM $Mg^{2+}$, 2.5mM glutathione, 10% glycerol, pH 7.4) by centrifugation at 4,500 x g for three minutes. Subsequently, to wash away excess nucleotides 100 µl of GXP:H-Ras complex was mixed with 500 µL Ligand Buffer in each concentrator, then spun at 4,500 x g for 5 min thereby returning the complex volume back down to 100 µl. This step was repeated four more times for a total $(1/5)^5$ = 3000-fold wash of the complexes with ligand buffer. Next the washed GXP:H-Ras complexes were transferred to ultracentrifuge tubes and were spun at 80,000 rpm for 10 min at 4º C in a Beckman TL-100 tabletop ultracentrifuge to eliminate aggregates. The UV deconvolution protocol was performed as previously described[2,3] to determine the total concentration of Ras protein. This total concentration was converted to the concentration of active Ras by multiplying with the fraction of active Ras, as determined by above HPLC quantification. The concentration of active Ras was adjusted to 100 µM for the working Ras stock in Ligand Buffer. The RBD construct MBP-RBD-GFP was thawed on ice and adjusted to a concentration of 200 nM for the 2X RBD stock in Target Buffer (25 mM HEPES, 140 mM KCl, 15 mM NaCl, 1 mM $Mg^{2+}$, 2.5 mM glutathione, 1 mg/mL BSA, 1% Tween 20).

Each MST titration series was generated and subjected to MST analysis as follows[2]. Sixteen Ras dilutions of the working Ras stock were carried out to create a set of 2X Ras stocks in Ligand Buffer, then each was mixed 1:1 with the 2X RBD stock in Target Buffer to yield three triplicate samples. The resulting three sets of 16 samples possessed the same, fixed RBD concentration (100 nM) and varying active Ras concentrations (50 µM, 50 µM, 40 µM, 35 µM, 30 µM, 25 µM, 20 µM, 16 µM, 12 µM, 8 µM, 4 µM, 2 µM, 1 µM, 0.5 µM, 0 µM, 0 µM). Approximately 7.5 µL of each sample was loaded into an MST capillary for measurement, yielding three triplicate titration series. Each titration was then measured separately in the Nanotemper Monolith NT.115 instrument controlled by MO.Control software. Settings included blue excitation light source at 10% intensity, high IR laser power (yielding a 6°C temperature increase above ambient at the center of the thermal gradient), sample chamber controlled at 22 °C, and capture of the MST signal 1.5 s after activation of the IR laser. Raw data, including the MST signal measured for each sample in a given titration, was exported using MO.Analysis software. All titrations were carried out in 25 mM HEPES, 140 mM KCl, 15 mM NaCl, 1 mM $Mg^{2+}$, 2.5 mM glutathione, 0.5 mg/mL BSA, 0.5% Tween 20. The detergent Tween 20 is a standard component in MST titrations (see https://nanotempertech.com/blog/top-6-ways-to-optimize-your-mst-assay/) and serves to prevent protein binding to capillary walls. We found that the detergent was also useful in preventing protein aggregation in our titrations, especially at high Ras concentrations. The present studies utilized our published MST procedure that has been extensively validated, including the use of 0.5% Tween 20 in titrations[2].

In each experiment, side-by-side triplicate titrations were carried out for the experimental H-Ras mutant and the control mutant H-Ras Q25L, which was used as an internal standard in MST measurements to confirm that the instrument and experiment were nominal, and to quantify the maximal MST binding signal (Bmax) for the day[2]. MST titration data was analyzed as previously described[2]. The MST signals obtained for the two negative control (0 µM Ras) samples in the same titration were averaged and the resulting baseline value was subtracted from the MST signals of all sixteen samples of that titration, yielding their change in value from the average “zero-point”. The resulting three baseline-corrected triplicate titrations were averaged to generate a mean active Ras titration. To account for any underlying trends in the instrument or sample background, “blank” runs (6 triplicate titrations) were performed in which the RBD construct was titrated with Ras-free ligand buffer, then the 18 blank runs were baseline corrected and averaged to give a standard, average blank titration. The resulting standard blank titration was subtracted from each mean Ras titration to yield the buffer-corrected, mean active Ras titration.

Once the buffer-corrected, mean Q25L active Ras titration was obtained, Prism software (GraphPad, San Diego, CA) was used to best-fit the binding equation $Y = (Bmax * X)/(X + K_D)$, where X is the active Ras concentration, by nonlinear regression. The resulting best-fit Q25L Bmax parameter represented the MST signal at full RBD saturation with bound Ras, and was found to be the same, within error, for Q25L and most other high affinity proteins (for exceptions see Table 1 in the text). Next, for all mean active Ras titrations measured on the same day, each mean data point was divided by the best-fit Q25L Bmax value, thereby converting each data point in the titration to a fractional RBD occupancy. Finally, the mean active Ras titration for each Ras protein was plotted as Fractional Occupancy of RBD against [Free Active HRas] (Figure 2), and the best fit $K_D$ value was determined by best fit of the binding equation $Y= [X]/([X] + K_D)$ via nonlinear regression where X is [Free Active HRas]. We set free [Free Active HRas] = [Total Active HRas] since the experimental conditions ensure these quantities differ by < 5% in all cases. Occasionally a high affinity mutant yielded a binding curve for which the asymptote was reproducibly greater or lesser than the value of 1.0 required for full occupancy of the single Ras binding site on the $PI3K\gamma_{RBD}$ construct. This feature arose when the mutant possessed a maximum MST signal larger or smaller than the Bmax measured for Q25L. For such a mutant the binding curve was best fit to the binding equation $Y = C * [X]/([X]+K_D)$. Then the data points for the mutant were individually divided by C to normalize the curve and plotted (Figure 2), and the normalization factor C presented in Table 1. The MST signal is determined by the effect of complex formation on multiple parameters including molecular mass, volume, shape, conformational dynamics, surface charge, hydration, and the quantum yield of GFP, thus it is not possible to extract useful information from differences in the normalization factor[1].

**Statistics.** To ensure the rigor and reproducibility of $K_D$ measurements, for each Ras protein at least 8 triplicate titrations were carried out on different days by at least 2 undergraduate Honors researchers. Moreover, the measurements for a given Ras protein employed at 2 separate protein preps. The resulting 8+ mean triplicate $K_D$ values were averaged to yield a global average $K_D \pm$ SEM, where the standard error of the mean was calculated for the total number of triplicates. Finally, statistical analyses were carried out using a two-tailed T-test to calculate p values to determine whether the $K_D$ value measured for WT H-Ras was significantly different from $K_D$ values measured for variants showing weaker binding (D33E, I36V, S39Y, R41L, and GDP-Ras) or tighter binding (Q25L, E37K, R41K and R41Q) or equivalent binding (G13R). T-tests were carried out using PRISM software (Graphpad, San Diego, CA) where $p < 0.05$ was required for significance.

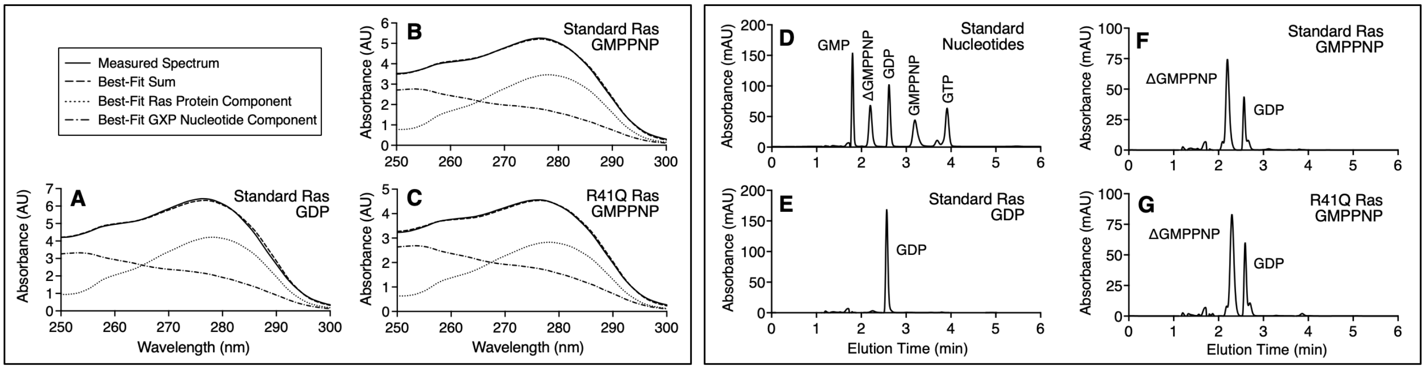


**Figure S1. Representative UV Deconvolution and HPLC Analyses Used to Quantify the Concentration and Nucleotide Loading of H-Ras Stocks for Binding Measurements.** Detailed procedures are published[3, 4] and provided above in Methods. **(A-C)** Examples of Nanodrop UV deconvolutions that resolve the UV spectrum of a given Ras stock solution into its protein and guanine nucleotide components and their concentrations. Each data set is an average of two triplicate measurements. **(D-G)** Examples of HPLC traces that quantify standard nucleotides (D), or nucleotides extracted from Ras proteins loaded with: (E) 100% GDP (only peak); (F) 70.8% ± 0.4% GMPPNP, remainder GDP; (G) 67.3% ± .2% GMPPNP, remainder GDP. The gentle nucleotide exchange procedure employed is optimized for retention of protein activity rather than full loading with the desired nucleotide, making it essential to quantify the loading percentage. Each trace is an average of triplicate measurements.

**Table S1. Effects of 20 Interfacial Ras Mutations on Ras Binding to $PI3K_{RBD}$ as Measured for the H-Ras:$PI3K\gamma_{RBD}$ and K-Ras:$PI3K\alpha_{RBD}$ Isoform Pairs**

| Ras Protein | Affinity for $PI3K_{RBD}$ Relative to WT | Number of COSMIC Cancer Database Samples Total / per Isoform[d] | Minimum Number of Base Changes[e] |
|---|---|---|---|
| | *Affinity Increase* | | |
| H-Ras R41Q[a] | 6.8-fold | 4 / HRas(4), NRas(0), KRas(0) | 1 |
| H-Ras E37K[a] | 5.2 | 2 / HRas(0), NRas(0), KRas(2) | 1 |
| **K-Ras Q25A[c]** | **4.1** | **0 / HRas(0), NRas(0), KRas(0)** | **2*** |
| **K-Ras E31D[c]** | **3.3** | **1 / HRas(1), NRas(0), KRas(0)** | **1** |
| H-Ras Q25L[a] | 2.4 | 1 / HRas(1), NRas(0), KRas(0) | 1 |
| H-Ras R41K[a] | 2.0 | 3 / HRas(0), NRas(1), KRas(2) | 1 |
| | *Similar to WT* | | |
| **K-Ras R41T[c]** | **1.4-fold** | **0 / HRas(0), NRas(0), KRas(0)** | **1** |
| **K-Ras H27Y[c]** | **1.2** | **2 / HRas(0), NRas(0), KRas(2)** | **1** |
| **K-Ras Y64F[c]** | **1.2** | **0 / HRas(0), NRas(0), KRas(0)** | **1** |
| H-Ras[a,b], **K-Ras[c]** | 1 (WT) | NA | NA |
| | *Affinity Decrease* | | |
| **K-Ras S39A[c]** | **0.30-fold** | **0 / HRas(0), NRas(0), KRas(0)** | **1** |
| **K-Ras D38A[c]** | **< 0.2** | **0 / HRas(0), NRas(0), KRas(0)** | **1** |
| H-Ras Y40C[b] | < 0.2 | 0 / HRas(0), NRas(0), KRas(0) | 1 |
| H-Ras R41L[a] | < 0.2 | 0 / Blocks v-Ras Transformation[8] | 1 |
| H-Ras I36V[a] | < 0.2 | 1 / HRas(1), NRas(0), KRas(0) | 1 |
| H-Ras S39Y[a] | < 0.2 | 2 / HRas(0), NRas(0), KRas(2) | 1 |
| H-Ras D38E[b] | < 0.2 | 1 / HRas(1), NRas(0), KRas(0) | 1 |
| H-Ras D33E[a] | < 0.2 | 16 / HRas(0), NRas(1), KRas(15) | 1 |
| **K-Ras I36A[c]** | **< 0.2** | **0 / HRas(0), NRas(0), KRas(0)** | **2*** |
| **K-Ras Y40A[c]** | **< 0.2** | **0 / HRas(0), NRas(0), KRas(0)** | **2*** |
| **K-Ras Y64A[c]** | **< 0.2** | **0 / HRas(0), NRas(0), KRas(0)** | **2*** |

[a] Affinity of the indicated H-Ras variant for $PI3K\gamma_{RBD}$ was measured using published procedures[2] detailed above in Methods, then was ratioed to the measured affinity for WT H-Ras binding to $PI3K\gamma_{RBD}$.

[b] Same, measured by Fleming et al[2].

[c] Affinity of the indicated K-Ras variant for $PI3K\alpha_{RBD}$ was measured by Czyzyk et al[6] and ratioed to the measured affinity for WT K-Ras binding to $PI3K\alpha_{RBD}$.

[d] Total number of tumor samples associated with the indicated mutation in H-Ras, N-Ras or K-Ras as tabulated by the COSMIC cancer database[7].

[e] Minimum number of base changes needed to introduce the indicated mutation.

*Note changes of >1 base are unlikely in the COSMIC somatic mutation database.

**CONTRIBUTIONS:**

This undergraduate research project was carried out by UC Boulder Honors students in the Falke laboratory, including all microscale thermophoresis (MST) titrations, as follows: EHM (MST, Figure 3C-E, Table 1). KCB (MST, Figure 2B-D, supported in part by CU Boulder UROP Award), KHS (MST), IRF (MST, supported in part by Beckman Scholar Award), CDT (MST, supported in part by Beckman Scholar Award), AMH (MST, supported in part by CU Wuttke/Beckman Biomedical Sciences Research Award), AL (MST). Support was provided by senior lab associates: LL (HPLC) JRA (HPLC), JPH (HPLC, Figures 1C, 3A), AHE (MST Training and Expertise). PI was JJF (Funding, Manuscript, Final Figures and Tables).
